# SPA-C: an hybrid tool to accurately scaffold genomes using Hi-C and Deep-Learning

**DOI:** 10.64898/2026.06.16.732382

**Authors:** Alexis Mergez, Raphaël Mourad, Guillermina Hernandez-Raquet, Matthias Zytnicki

## Abstract

Genome assembly is a computational pipeline designed to reconstruct chromosomes from small sequencing reads. Following their assembly, contiguous sequences (contigs) are arranged into chromosome-long sequences during scaffolding. Hi-C, a long-range linkage information between regions of the genome widely used in recent large sequencing projects, is often required to correctly order contigs. Several tools have been developed to automate this task following either statistical or deep-learning approaches. Statistical approaches summarise 2D Hi-C matrices into contact densities across sequences, thus ignoring informative visual patterns. The sole existing deep-learning tool uses a transformer-based computer vision model to correct the assembly. It has been trained on several species and uses Hi-C matrices directly. Yet it comes as a supplementary step in the scaffolding process, introducing extra computation time, and has been trained on a dataset that might contain labelling errors, which could provide sub-optimal results. We propose *SPA-C*, an hybrid pipeline combining the strengths of both approaches. Linkage prediction is handled with a frugal CNN-based model and a graph-solving algorithm is used to generate the scaffolds. Through our input’s design, the model is able to both correct errors within assemblies and link contigs, leveraging small, local Hi-C contact matrices. We handled low-complexity regions that might induce erroneous predictions using an external tool, improving the overall accuracy of generated assemblies. On a benchmark of six various genomes and four standard metrics, *SPA-C* outperformed four out of four state-of-the-art methods while achieving comparable start-to-end computation time. Python and Bash scripts are available on GitHub (github.com/SPA-C/SPA-C.git) and Zenodo (10.5281/zenodo.19000361).

## 1 Introduction

Genome assemblies, as fundamental objects of genomics, are used in genetics, evolutionary biology, and pangenomics. Genome quality is essential as it directly impacts downstream applications. For instance, CHM13 [Rhie et al., 2023], the first complete, accurate and gapless (telomere-to-telomere, T2T) human genome assembly, added 140 protein-coding genes and removed falsely duplicated regions compared to the GRCh38 reference assembly [Nurk et al., 2022]. It also improved structural variants detection accuracy which contributed to the discovery of 4 inversions linked to human genetic diseases [Rocha et al., 2024].

*De novo* genome assembly is a two-step computational process to reconstruct a genome from fragmented sequences called *reads*: contig assembly and scaffolding. As the complexity of this task lies in the quantity and length of repetitive regions, the advent of PacBio HiFi (10 − 20 kb, ≤ 0.5% error rate) accurate long-reads and ONT ultra-long reads (≥ 100 kb, 1 − 2% error rate) enabled researchers to elucidate most of these regions [Li and Durbin, 2024]. During contig assembly, contiguous sequences are built using overlaps between reads [Cheng et al., 2021], generating an initial assembly. Even using long-reads, contigs usually do not reach the full length of chromosomes [Schell et al., 2025].

While the Hi-C sequencing protocol was initially developed to study the 3D conformation of the genome, it also captures the 1D proximity between sequences [Lieberman-Aiden et al., 2009]. This information is useful to both scaffold contigs into chromosomes [Burton et al., 2013] and phase assemblies into haplotypes [Cheng et al., 2021]. Several tools have been developed to automate Hi-C-based scaffolding using statistical or deep-learning approaches. Both rely on an iterative process involving assembly error correction by breaking contigs and scaffolding. Statistical approaches such as *YaHS* [Zhou et al., 2023], *Pin hic* [Guan et al., 2021] *and 3DDNA* [Dudchenko et al., 2017], use contact expectation models that summarise 2D contact matrices into contact density, computed at each *locus*. While offering a good proxy in most cases, they discard the informative visual patterns that could help detect smaller or more complex assembly errors [Howe et al., 2021]. The novel deep-learning approach applied in *AutoHic* [Jiang et al., 2024], instead relies on a Swin transformer model to directly detects erroneous patterns from contact matrices. It addresses the density issue, yet the model was trained on non-T2T assemblies and pseudo-labelled using a 90% accurate deep-learning model [Jiang et al., 2024]. Both could introduce labelling errors and biases as assemblies might contains misjoins being misclassified. *AutoHic* also relies on *3D-DNA* outputs, functioning as an additional step in the scaffolding process, increasing the overall computation time.

Here we propose *SPA-C*, a hybrid Hi-C scaffolder that combines the strengths of both statistical and deep-learning approaches. Contig breaking and scaffolding are both predicted using a single frugal ResNeXt [Xie et al., 2017] model, achieved through a specific design of its inputs, and trained on a curated dataset based on the CHM13 T2T assembly. Similar to *AutoHic*, the model is able to use the 2D patterns of the Hi-C contact matrix while achieving faster inference. After a first error screening step, linkage predictions are aggregated into a scaffolding graph that is later solved using the *YaHS* scaffolding algorithm. *Longdust* [Li and Li, 2025] was used to improve the tool’s overall accuracy by discarding predictions of the model in low complexity regions that tend to generate erroneous patterns. We trained the model on synthetic errors with a fixed resolution and found it was able to generalise to other species, other resolutions, and handle missing information. Compared to previous scaffolding tools, we found that SPA-C reconstructed curated genomes of six different species more accurately and in an equivalent time as YaHS, one of the fastest tool.

## 2 Material and Methods

### 2.1 Pipeline overview

We defined “misjoins” as an erroneous link between two non-contiguous sequences and “joins” as a correct connection between contigs. Following the creation of Hi-C contact matrices, misjoins that might have been introduced during contig assembly are detected using a sliding window along contig’s intra matrix diagonals (see Fig.S1). Joins between correct contigs are then predicted using the same model and summarised into a scaffolding graph (GFA). This graph is finally solved using a reimplementation of *YaHS* scaffolding rules, yielding a final FASTA file containing scaffolds (see Fig.S2).

### 2.2 Hi-C matrix generation

Hi-C contact matrices were generated using a custom workflow (see Fig.S3). *Juicer* [Durand et al., 2016] was used to align Hi-C reads (FASTQ) to contig-scale assemblies (FASTA) and pre-process alignments, yielding a de-duplicated BAM alignment file. *HicExplorer* [Wolff et al., 2020] was then used to generate MCOOL files from the deduplicated alignments (see Supp. Methods). All other scaffolders also rely on *Juicer*. We favored the MCOOL format over the HIC format, as it supports faster data processing.

### 2.3 Contig breaking

Due to the layout of samples seen during training (see Fig.S4), the model predicts a joining score (pseudo-probability) between two consecutive regions of a sequence. To detect misjoins within input contigs, the model compute scores using a sliding-window along the diagonal of its Hi-C matrix. As both forward and reverse matrices are independently used for prediction, scores are simply averaged. Thus, we obtain a prediction curve along contigs varying between 0 and 1, where 0 indicates a misjoin. As matrices overlap each other (see Fig.S1), predictions also overlap and a misjoin can be seen in several matrices surrounding its actual bin position. Therefore, the prediction curve does not indicate a precise bin position but rather a range of positions around the misjoin, which we later referred as “chimeric regions” (see Fig.3). We used a threshold of 0.05 to classify misjoins and we used the middle bin position of chimeric regions ≤ 19 bins as the misjoin position (Fig. 3A-C). Longer chimeric regions were not split and instead treated as contig, as they might contain several misjoins that cannot be accurately located.

Low complexity regions tend to induce misleading patterns in the contact matrix which are then misclassified by the model (see Fig. 3D). *Longdust* [Li and Li, 2025] was used to detect such regions from the sequence itself and improve the misjoin detection overall accuracy. Low complexity regions of size greater than 2 × bin resolution (i.e. 10 kb regions by default) were not classified as misjoins, even if the model suggested the contrary.

### 2.4 Scaffolding

Contigs that have been screened for misjoins and corrected if needed, are then scaffolded into chromosomes. To achieve this, we reused the same model (without retraining) to predicts linkage pseudo-probability between each pair of corrected contig’s ends where a score of 1 indicates a join. We computed scores using both orders (A-B, B-A) and all four orientations: *start-start, start-end, end-start* and *end-end*. Equivalent links were averaged between contig pairs. For instance, link A:end-B:start is equivalent to link B:start-A:end, but both matrices are separated by a 180° rotation. Note that compared to the 2.3 section, no sliding-window is used, therefore, two matrices are used to assess the “joining” score for a given orientation. By default, we used two bin resolution (5 kb and 25 kb) to predict linkage scores and then we averaged equivalent link scores, yielding a single Graphical Fragment Assembly file (GFA). In this oriented graph, nodes are corrected contigs sequences and edges are weighted using predictions of the model. This graph is passed to a re-implementation of *YaHS* [Zhou et al., 2023] scaffolding algorithm only, modified to support GFA-based inputs. For each contig’s end, the algorithm chooses the best target node and orientation from all possible links using edges’ weights. We chose this algorithm as it makes a choice if a score is clearly better than other possibilities, and as it handles complex topologies arising from repeats or small contigs. This limited the number of misassemblies overall.

### 2.5 Model

#### 2.5.1 Architecture

The *SPA-C* model is based on the ResNeXt architecture [Xie et al., 2017], that we significantly downsized to 42,000 parameters. As our input size (20 *×* 20) is smaller than the original input size of ResNeXt, we removed several blocks of convolution (see Tab.S1 for a comparison). The model features an encoder (see Fig. 1) and a classifier. The encoder takes normalised Hi-C matrices and extracts meaningful features into an embedding of size 64. This embedding is passed to the classifier which predicts its label as pseudo-probability: 0 for a misjoin and 1 for a join. Classic convolutions were replaced with partial convolutions [Liu et al., 2018] to allow for masked samples.

**Figure 1:**
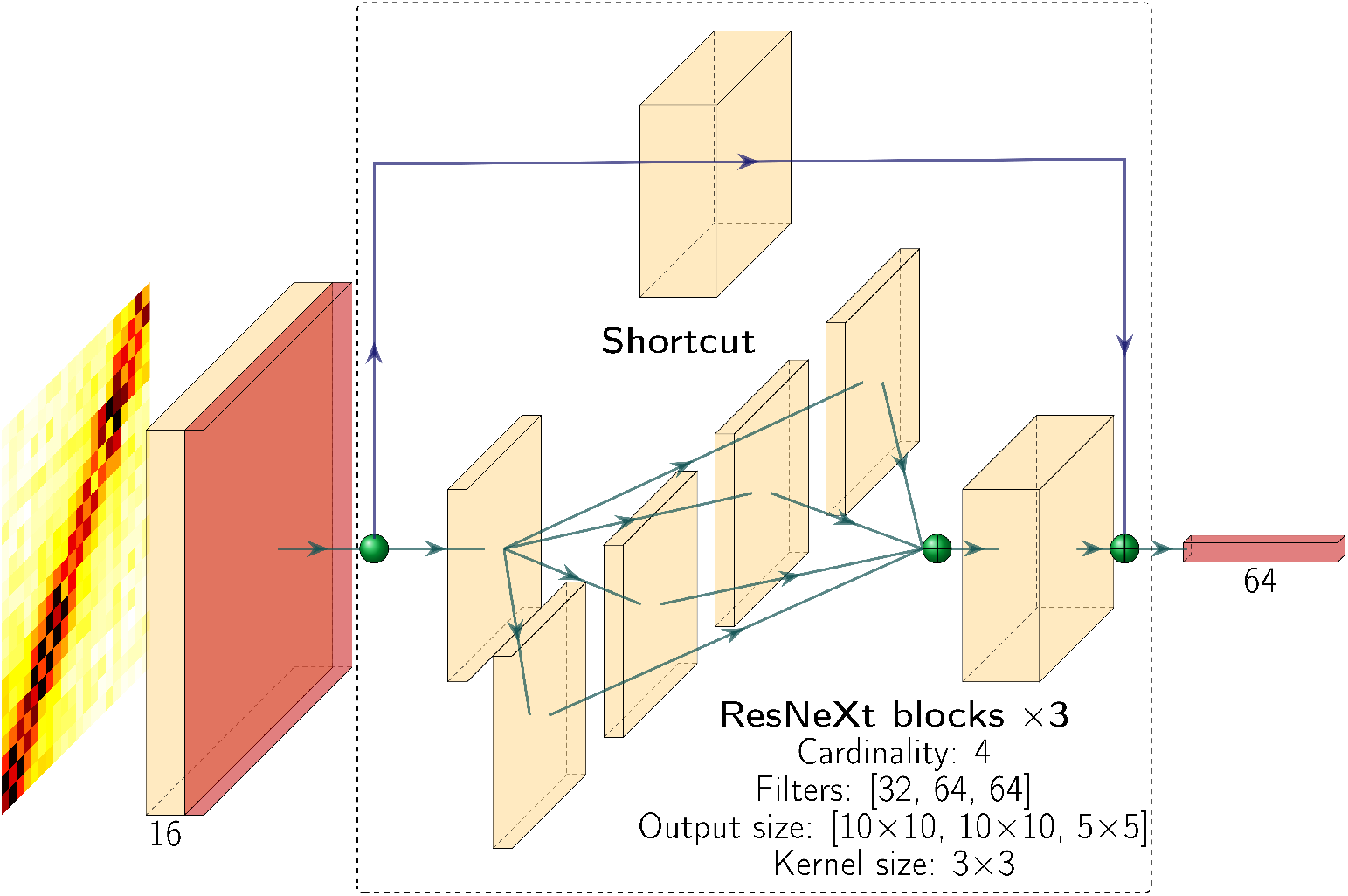
Encoder architecture of *SPA-C*, based on ResNeXt. A first convolution reduces the size of the input down to 10 *×* 10. This is followed by three ResNeXt blocks (cardinality of 4): two increasing the number of channels (8 to 32, 32 to 64), and one reducing the dimension (10 to 5). A final average pool layer brings the embedding to a size of 64.

#### 2.5.2 Datasets

In this paper, we distinguished “intra” matrices containing contacts between regions of the same contig or chromosome, from “inter” matrices representing contacts between 2 distinct contigs. We designed the model’s input using the arrangement of “intra” matrices (see Fig.S4): the diagonal is visible and anti-quadrants contains the contacts between the two regions. Therefore, each “inter” sample is equivalent to an “intra” sample of a contig that contains a misjoin between sequences from two different chromosomes (Fig. 3A). Using this input arrangement, we hypothesised that a model that classifies “misjoins” and “joins” would be able to both detect misjoins in input contigs and predict linkage between contigs.

To prevent any labelling errors, we used gold standard human T2T assemblies CHM13v2 [Rhie et al., 2023] and the maternal haplotype of HG002v1.1 [Hansen et al., 2025] to respectively train and test the model (see Tab.S2 for accessions). The *cooler* Python library [Abdennur and Mirny, 2020] was used to extract 20 *×* 20 matrices from MCOOL Hi-C matrices, which are used as input for the model. Join (correct) samples were found using a sliding-window mechanism along correctly assembled chromosomes (see Fig.S1A) and were labeled as “1.” Since accurately detecting errors in an assembly is a challenging task without a ground truth, we instead chose to simulate assembly errors using the same T2T assemblies. This allowed us to both mimic and generate many samples for all possible type of misassemblies: inversions, inter-chromosomal mijoins and deletions. Chimeric contigs or chimera (contigs containing misjoins) were designed as 200 kb long sequences, divided into two equally-sized regions. Chimera of each type of misassembly were generated by randomly sampling regions without replacement. Using distant regions of the same chromosome with gap length varying between 100 kb and 5 Mb, we simulated 15,179 deletions. Fig.S5 reports the distribution of gap length. Inversions were generated using two contiguous regions and reversing only one, yielding 13,454 chimera. 15,238 inter-chromosomal misjoins were generated by sampling regions from different chromosomes. Misjoin samples were found using a sliding-window along chimeras, where the breakpoint is located at *±*3 bins from the center of the sample, and labeled as “0” (see Fig.S1B).

In addition to synthetic contigs, we generated misjoin samples from inter-chromosomal regions of correct assemblies. Matrices with fewer counts were sub-sampled in favor of samples with more complex patterns (see Fig.S6C).

#### 2.5.3 Training

The training dataset contained 2.35M samples with 1.22M join samples, representing 52% of the total. Misjoin samples included 728,055 inter-chromosomal misjoins, 212,506 deletions and 188,356 inversions. The dataset was split into three subsets using a stratified group split: *training* (90%), *validation* (5%) and *test* (5%), with non-overlapping groups. For instance, join samples from a given chromosome were grouped in the same subset, preventing data leakage. The model was trained using the *training* subset and validated with the *validation* subset to avoid overfitting at each epoch. Samples were randomly masked to simulate missing bins (see Fig.S7). We used the Binary Cross Entropy loss with logits and AdamW [Loshchilov and Hutter, 2019] optimiser combined with a cosine annealing scheduler. The scheduler decreases the learning rate gradually to help the model converge, starting at 0.001 and decreasing gradually in ten epochs. After training, we selected the epoch 2 weights’ as the model achieved its lowest validation loss (0.01147) (see Fig.S8). Model performances were then assessed on the test subset.

#### 2.5.4 Normalisation

Since Hi-C sequencing projects and biological variability may lead to different Hi-C coverage between genomes, we normalised model’s inputs. Inspired by the colormap scaling used in *AutoHic* [Jiang et al., 2024], *we used a min-max scaling, applied on the matrix values itself. As maximum value, we used the 95th quantile computed from contacts located between the* −10 and +10 subdiagonals (Fig.S9) of chromosomes or contig “intra” matrices. After scaling, values were clipped between 0 and 1. Since high-resolution Hi-C contact matrices tend to be highly sparse, this computation method emphasises low-value counts that might otherwise be reduced to values close to zero.

### 2.6 Benchmark

#### 2.6.1 Data

To compare our approach to other Hi-C scaffolding tools, we used high-quality, curated genome assemblies from The Darwin Tree of Life (DToL) project [The Darwin Tree of Life Project Consortium, 2022]: *Arabidopsis thaliana, Drosophila histrio, Astur gentilis, Vulpes vulpes* and *Balaenoptera physalus* [Christenhusz et al., 2023, Obbard et al., 2024, August et al., 2022, Lopez Colom et al., 2025, Davison et al., 2026] (see Tab.S2 for accessions). We split each assembly at every scaffolding gaps back into contigs. Since the HG002v1.1 assembly is gapless, we instead split chromosomes of the maternal haplotype into 10 Mb long contigs. It should be noted that YaHS was used in all DToL genome assemblies used in this paper and that results may be favoring this tool.

#### 2.6.2 Metrics

Like [Zhou et al., 2023, Guan et al., 2021, Dudchenko et al., 2017], we assessed scaffolding performances by comparing NGA50, NGA90, auNGA and the number of misassemblies as reported by *Quast* [Mikheenko et al., 2023]. The NGAx summarises the similarity between query assemblies and a reference assembly by combining the Nx metric and the number of misassemblies [Gurevich et al., 2013]. Briefly, assembly contigs are aligned to a reference, broken at each misassembly breakpoint, and sorted by decreasing length. For a given x, the NGAx reports the length of the non-erroneous contig such that x% of the reference length is achieved. The higher the value, the closer the assembly is to the reference. These metrics are standard when comparing genome assembly tools.

## 3 Results

### 3.1 Classification

We assessed the model’s classification performance using the test subset of CHM13v2 (Fig.S10) and the HG002v1.1 maternal haplotype dataset. We built this second dataset using the same method as the training dataset. It contains 2.3 M samples (53% joins) and allowed for an evaluation on different Hi-C data. The model achieved AUC-ROCs of 1 and 1, and AUPRs of 0.999 and 0.998 on the test subset and the HG002 dataset respectively (Fig. 2). We subsequently chose a threshold of 0.05 as it did not impact classification performances and it made our approach more conservative, under the assumption that contigs are rarely containing misjoins.

**Figure 2:**
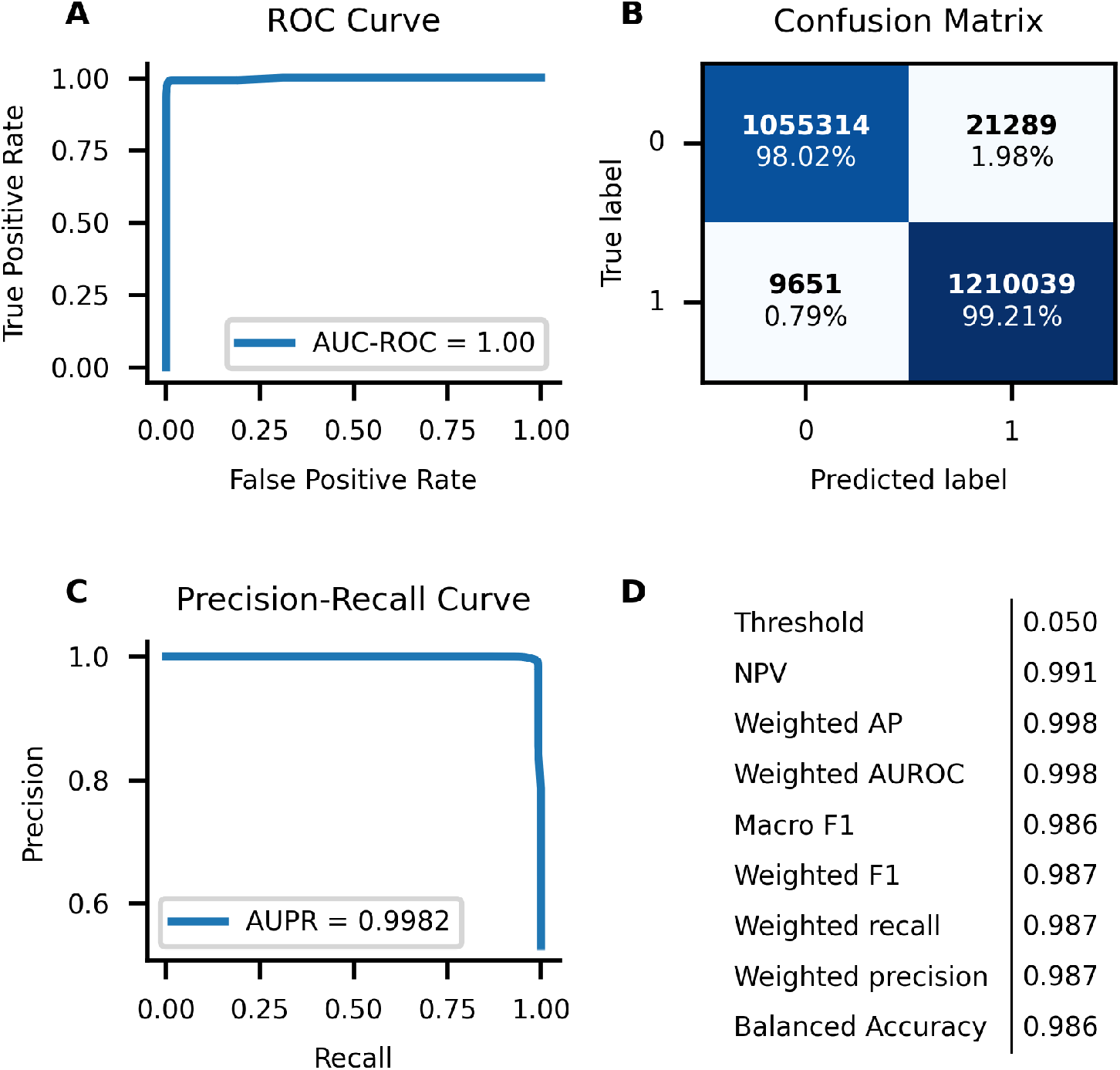
Classification performance of *SPA-C* model on the HG002 maternal haplotype dataset. (A) ROC Curve and AUC-ROC. (B) Confusion matrix using a threshold of 0.05. Label 0 corresponds to examples of assembly error. (C) PR Curve and AUPR. (D) Several classification performance metrics using a threshold of 0.05.

Figure 3 shows some examples of assembly errors and their associated prediction curve. In Fig.3A-C, the model accurately predicts the breakpoint position (see section 2.3). Fig.3D shows a low complexity region (predicted using *Longdust*) that has been misclassified by the model, but will be discarded. To further assess the impact of *Longdust*, we compared assembly contiguity in Tab.S3, using the same genomes as in section 3.2. Although its impact vary between datasets, *Longdust* improved the scaffolding in every metrics, allowing a seven fold increase of the NGA90 on the *A. thaliana* dataset.

**Figure 3:**
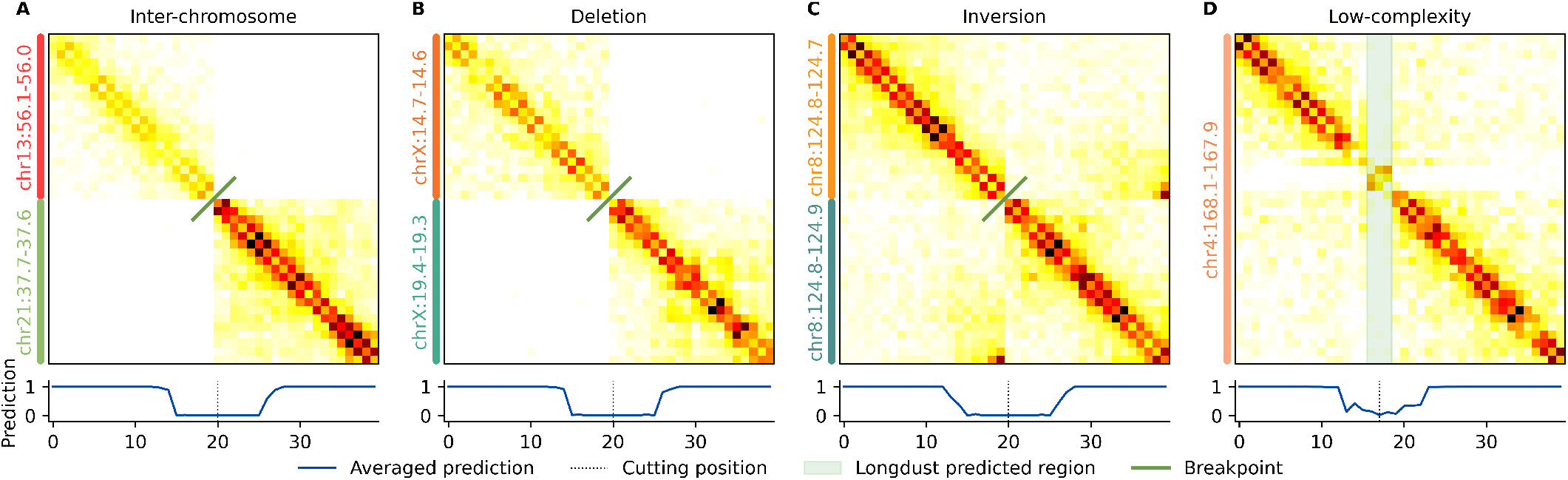
Several assembly error patterns and their associated predictions. The average prediction curve vary between 0 (“misjoin”) and 1 (“join”). Examples from the HG002 maternal haplotype dataset are shown using a 5 kb bin resolution. (A-C) Examples from synthetic chimeric contigs. Using our method, the contig is split at the misjoin position. The actual misjoin location is shown using the green line. (D) Example from a correctly assembled chromosome, in which the low complexity region was predicted using *Longdust*. The regions is misclassified by the model, but the contig will not be split.

We evaluated the model on other bin resolutions (see Tab.S4). Using longer chimeras (4 Mb) instead of the 200 kb detailled in section 2.5.2, the model performances remained excellent with an AUPR and an macro-F1 greater than 0.996 and 0.9861 respectively. Multi-resolution prediction is often used to capture longer-range proximity information and improve scaffolding [Zhou et al., 2023, Guan et al., 2021, Dudchenko et al., 2017]. We tested this by comparing our two resolution approach to a single resolution version in Tab.S5. Overall, multi-resolution improved genome contiguity.

### 3.2 Benchmarking on several species

Using six high-quality assemblies (see 2.6.1), we compared Hi-C scaffolding tools and summarised the results in Tab.1. CPU time and all *Quast* metrics are reported in Tab.S6 and Tab.S7 respectively. *SPA-C* achieved the best overall rank (1.62), followed by *YaHS* and *Pin Hic* (2.17 and 2.42 respectively). *3DDNA* and *AutoHic* were systematically last, achieving 3.93 and 4.75 respectively. In our testings, *AutoHic* yielded scaffolds containing ≤ 1% of the reference total base and was subsequently ranked last for the corresponding genomes. *SPA-C* ranked first on four out of six genomes, with one *ex æquo* with *YaHS. YaHS* ranked first on *V. vulpes* and *Pin hic* ranked first on *A. gentilis*. Except for *Pin hic, SPA-C* generated less misassemblies when compared to other tools, while achieving similar or more contiguous scaffolds.

**Table 1:** Metrics on several high-quality, manually curated genomes. NGA associated values and the number of misassemblies were computed using *Quast*. For each genome, the average rank of each tool was computed using its rankings on metrics and reported in “Dataset rank”. A global rank was also computed for each tool by averaging its rankings on all datasets.

|  | Dataset | # misassemblies | NGA50<sup>1</sup> | NGA90<sup>1</sup> | auNGA<sup>1</sup> | CPU time (h) | Dataset rank<sup>2</sup> | Global rank<sup>3</sup> |
| --- | --- | --- | --- | --- | --- | --- | --- | --- |
| YaHS | *A. gentilis* | 303 | 24.69 | 2.19 | 25.67 | 167 | 2.50 | 2.17 |
|  | *A. thaliana* | 22 | 23.05 | 11.19 | 23.77 | 310 | 2.25 |  |
|  | *B. physalus* | 907 | 20.88 | 0.40 | 22.67 | 842 | 2.75 |  |
|  | *D. histrio* | 61 | **28.85** | **0.81** | **21.78** | 226 | **1.50** |  |
|  | HG002 | 7 | 133.58 | 66.07 | 132.45 | 504 | 2.50 |  |
|  | *V. vulpes* | 277 | **55.41** | **12.96** | **60.09** | 476 | **1.50** |  |
| Pin_hic | *A. gentilis* | **23** | 29.66 | **3.00** | **28.12** | 169 | **1.00** | 2.42 |
|  | *A. thaliana* | 10 | 13.13 | 4.80 | 12.80 | 212 | 2.75 |  |
|  | *B. physalus* | **119** | 15.41 | **0.48** | 17.96 | 854 | 2.00 |  |
|  | *D. histrio* | 62 | 8.07 | 0.71 | 11.88 | 231 | 3.75 |  |
|  | HG002 | 4 | 135.24 | 50.22 | 131.44 | 513 | 2.50 |  |
|  | *V. vulpes* | **176** | 18.26 | 4.23 | 21.44 | 520 | 2.50 |  |
| 3D-DNA | *A. gentilis* | 736 | 4.62 | 0.42 | 6.25 | 217 | 4.00 | 3.93 |
|  | *A. thaliana* | 350 | 0.40 | 0.07 | 0.50 | 232 | 4.00 |  |
|  | *B. physalus* | 728 | 2.11 | 0.26 | 2.82 | 1159 | 3.75 |  |
|  | *D. histrio* | 138 | 14.31 | 0.56 | 16.27 | 246 | 3.50 |  |
|  | HG002 | 2158 | 2.55 | 0.33 | 6.27 | 770 | 4.30 |  |
|  | *V. vulpes* | 553 | 16.06 | 3.62 | 20.28 | 1321 | 4.00 |  |
| AutoHic | *A. gentilis* | 0 | N.A.<sup>4</sup> | N.A.<sup>4</sup> | 0.03 | 319 | 5.00 | 4.75 |
|  | *A. thaliana* | 354 | 0.40 | 0.07 | 0.50 | 239 | 4.50 |  |
|  | *B. physalus* | 0 | N.A.<sup>4</sup> | N.A.<sup>4</sup> | 0.00 | 1159 | 5.00 |  |
|  | *D. histrio* | 0 | N.A.<sup>4</sup> | N.A.<sup>4</sup> | 0.02 | 251 | 5.00 |  |
|  | HG002 | 2158 | 2.55 | 0.33 | 6.27 | 935 | 4.00 |  |
|  | *V. vulpes* | 1 | N.A.<sup>4</sup> | N.A.<sup>4</sup> | 0.00 | 1321 | 5.00 |  |
| SPA-C | *A. gentilis* | 124 | 21.43 | 2.61 | 22.52 | 175 | 2.50 | 1.62 |
|  | *A. thaliana* | **1** | **27.69** | **15.78** | **25.19** | 218 | **1.00** |  |
|  | *B. physalus* | 468 | **30.05** | 0.35 | **34.86** | 944 | **1.75** |  |
|  | *D. histrio* | **32** | 23.34 | **0.81** | 20.23 | 236 | **1.50** |  |
|  | HG002 | **3** | **154.34** | **78.91** | **153.69** | 541 | **1.00** |  |
|  | *V. vulpes* | 193 | 35.80 | 7.79 | 39.96 | 514 | 2.00 |  |

<sup>1</sup> Expressed in Mbp.
<sup>2</sup> Average rank based on ranks on NGA50, NGA90, auNGA and number of misassemblies.
<sup>3</sup> Tool’s average rank across datasets.
<sup>4</sup> For some genomes, AutoHic’s output contained $\leq 1\%$ of the reference total length and was rank last.

When investigating the results on *V. vulpes*, we found that *SPA-C* arranged unplaced contigs from the DToL assembly into chromosome-level scaffolds. We assessed the correctness of insertions by comparing the contact distribution between adjacent matrices with a Wilcoxon test using the “greater” alternative. For instance, if a contig *C* is placed between contig *A* and *B*, we compared inter contacts between *A*—*C* and *C* —*B* with inter contacts *A*—*B*. Some examples and their associated p-values are reported in Fig. 4,S11. In each identified insertion, the p-value was significant.

**Figure 4:**
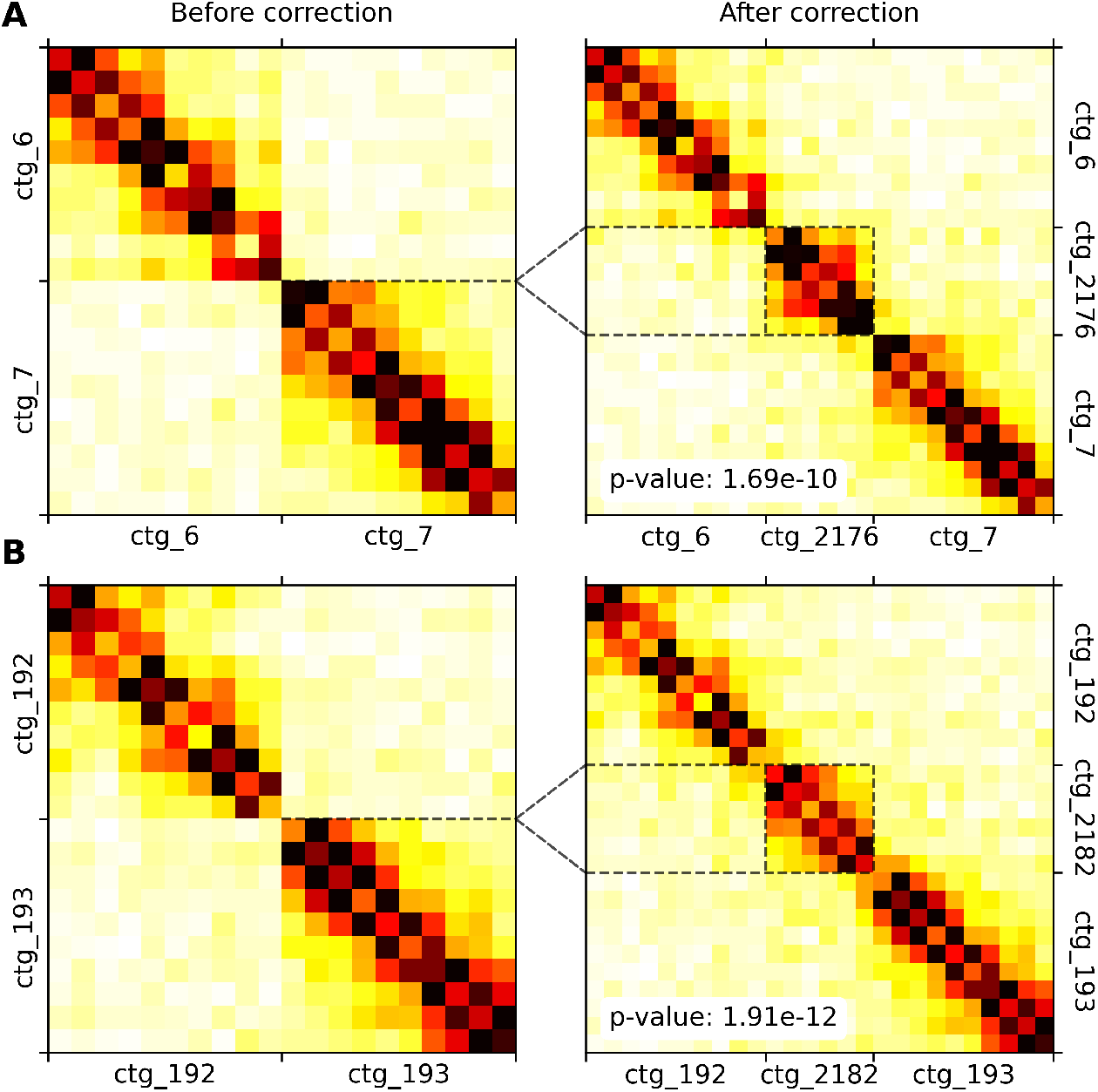
Improvements of the reference assembly made by *SPA-C* on the *V. vulpes* dataset. 5 kb bin resolution matrices are normalised. P-values are associated with the Wilcoxon “greater” alternative test between contact distribution before and after correction. Here, all p-values are significant, indicating that the distribution of contacts is greater after the insertion. Dotted lines show both the insertion position in the left matrix and the inserted contig’s boundaries in the right panel. (A) Example on Chromosome 1 of the reference assembly. (B) Example on Chromosome 2 of the reference assembly.

### 3.3 Sensitivity analysis

We tested the sensitivity of our approach to the Hi-C coverage by sub-sampling Hi-C contacts of the HG002 dataset from Tab.1. We found that a low to moderate under-sampling did not impact the model performances since matrices are normalised (Fig.5). But, for coverage below 0.03 reads per assembly base or 4.5X, the matrix has become too sparse for the model to make accurate predictions, leading to a decrease in NGA metrics.

**Figure 5:**
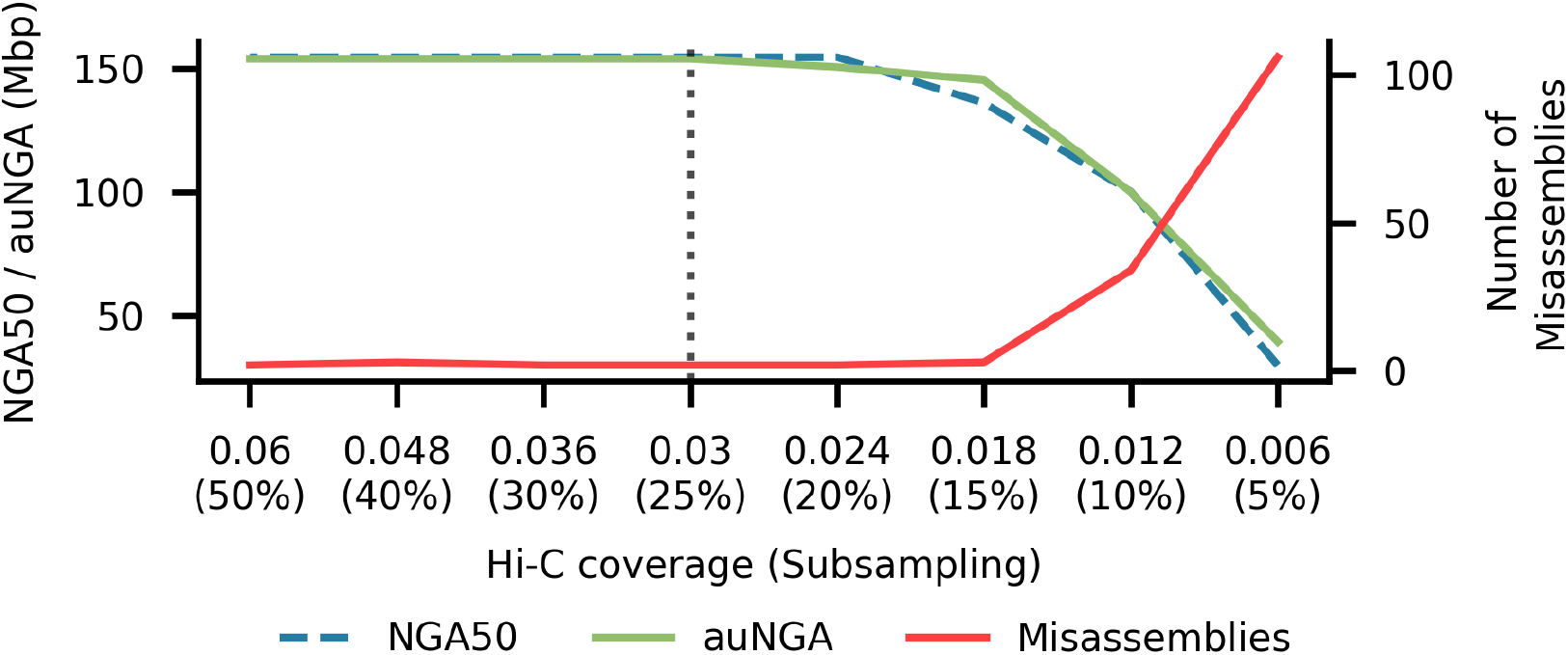
Sensitivity of *SPA-C* model to Hi-C coverage. We subsampled Hi-C reads of the 10 Mb contigs from the HG002v1.1 assembly dataset to simulate sparser Hi-C matrices. The coverage is given in Hi-C contacts per base. NGA50, auNGA and the number of misassemblies were computed using *Quast*.

Since smaller contigs might not fill the entire input size of the model (100 kb), we tested the model’s ability to handle missing bins (see Fig.S7) using partial convolutions [Liu et al., 2018]. To do so, we masked samples from the HG002 test dataset (Fig. 2) and found that the model was able to support up to 6 masked bins before the performance decreased. We compared masking to padding, in which missing values are replaced with neutral values (0 in our context), and showed that the former yields better results (see Fig.S7).

## 4 Discussion

In this paper, we proposed a frugal ResNeXt-based model, *SPA-C*, as part of a two-step hybrid approach to scaffold genome assemblies using Hi-C, focusing on achieving near-T2T quality and limiting manual curation efforts. On a benchmark of six various genomes and four standard metrics, *SPA-C* outperformed four out of four state-of-the-art methods while achieving comparable start-to-end computation time.

Compared to classical approaches relying on a iterative correction process, *SPA-C* uses a single-pass correction and joining approach, as well as a combination of tools to circumvent the limitations of Hi-C data, especially in low complexity regions. Instead of relying on a contact density (*YaHS, Pin hic, 3DDNA*), the model is able to capture spatial relationships directly from small-range, high-resolution, 2D Hi-C contact matrices. By carefully generating inputs, the model is used as a linkage predictor to both detect errors in input contigs and scaffold them.

It was trained using fixed-resolution matrices representing synthetic errors derived from the human CHM13v2 T2T assembly [Rhie et al., 2023] which minimised labelling errors, but might also have limited the diversity of samples. Despite this, it achieved excellent classification performances and was able to generalise to other bin resolutions, other species and handle missing information through masked inputs and partial convolutions. We selected a threshold of 0.05 as performances were robust and to minimise the risk of false negatives. We found that the model was able to precisely locate assembly errors, but showed some limitations with low complexity regions that tend to have uninformative Hi-C contact patterns. We solved it during error detection by discarding model’s prediction in *Longdust* predicted low complexity regions, improving error detection accuracy and final assembly quality. Finally, the model was able to accurately scaffold genomes with very low Hi-C coverage, far below the Earth Biogenome Project recommendations of 50X or 0.33 reads per bases [Lawniczak et al., 2022].

Following error detection and linkage prediction, a scaffolding graph is generated using all predictions and solved using *YaHS* algorithm which was re-implemented to support a GFA-based graph. As the model was trained to be sensitive to small structural errors (gaps, inversions), it very often favors a single orientation between contig pairs, allowing the algorithm to easily pick the correct orientation and order of contigs. The greater NGA metrics and greater number of contigs generally observed in our benchmark seems to indicate that our approach is more conservative compared to other scaffolders. When investigating *V. vulpes* dataset to understand *YaHS* performances, we found that *SPA-C* improved the quality of the curated assembly by properly placing left-out contigs into their chromosomes, but led to underestimated NGA metrics and inflated misassemblies counts. Since *YaHS* was used to generate the curated assembly, it could explain the better results it achieved compared to all other Hi-C scaffolders on this particular dataset. When predicting joins between low complexity regions (telomeres, centromeres for example), the model might be unable to both accurately predict the correct orientation and adjacent pairs of contigs. To avoid this issue, telomeric joins were removed using the same filter *YaHS* uses and multi-resolution prediction was implemented (Tab.S5) to capture longer range proximity information. Regarding *AutoHic*, we contacted its authors and they confirmed that we did not misused the tool.

There are some limitations to the approach. The model uses fixed size inputs, setting the minimum length of contigs to 50 kb in error detection and 25 kb in joining prediction (accounting for masked inputs). Therefore, smaller contigs are excluded of the pipeline but are still included in the final FASTA file. It should be noted that smaller contigs might also be difficult to arrange as Hi-C might not provide enough signal to predict their respective orientation. As we trained the model using human genome only, some genomes that induce specific Hi-C patterns might limit the scaffolding accuracy. *AutoHic* has been trained on several species using Hi-C data from the DNA Zoo library [Jiang et al., 2024] and does not have this limitation. Yet, we shown that our approach still outperforms theirs on diverse species.

*SPA-C* is a fast and accurate Hi-C scaffolder pipeline aimed towards near-T2T assembly. By limiting the number of misassemblies and improving scaffolding while achieving similar computation time compared to *YaHS*, manual curation tend to be limited to chromosome merging only. Our tool is compatible with data generated by large sequencing projects under the umbrella of the Earth Biogenome Project [Lawniczak et al., 2022], and could allow for significant time savings. Finally, our model was able to almost perfectly reconstruct a human T2T assembly, a critical task in the context of Human pangenomics which is aimed towards building several hundreds haplotypes large pangenome graphs, requiring the best assembly quality possible [Liao et al., 2023].

## 5 Competing interests

No competing interest is declared.

## 6 Author contributions statement

R.M., G.H-R., M.Z conceived the project. A.M. wrote the code, conducted the experiments and wrote the initial draft. A.M., R.M., G.H-R. and M.Z. reviewed the manuscript.

## 7 Acknowledgments

This study was funded through the “AAP Emergence 2024” program from the French Occitanie Region and the “Investissements d’Avenir” program of the French National Research Agency under the reference ANR-18-EURE-0021. We thank colleagues of the MIAT lab and especially Xian C. for her careful review of the manuscript. We are grateful to the Bioinfo-Genotoul platform (https://doi.org/10.15454/1.5572369328961167E12) for providing help, computing and storage resources.

## 8 Code and Data availability

All data used in this research are publicly available with accessions given in the main text and in Tab.S2. The code is available at github.com/SPA-C/SPA-C.

## Notes

https://github.com/SPA-C/SPA-C.git

https://doi.org/10.5281/zenodo.19000361

### Competing Interest Statement

The authors have declared no competing interest.

