## Supplementary Figures and Tables 1-6 for "SPA-C: an hybrid tool to accurately scaffold genomes using Hi-C and Deep-Learning"

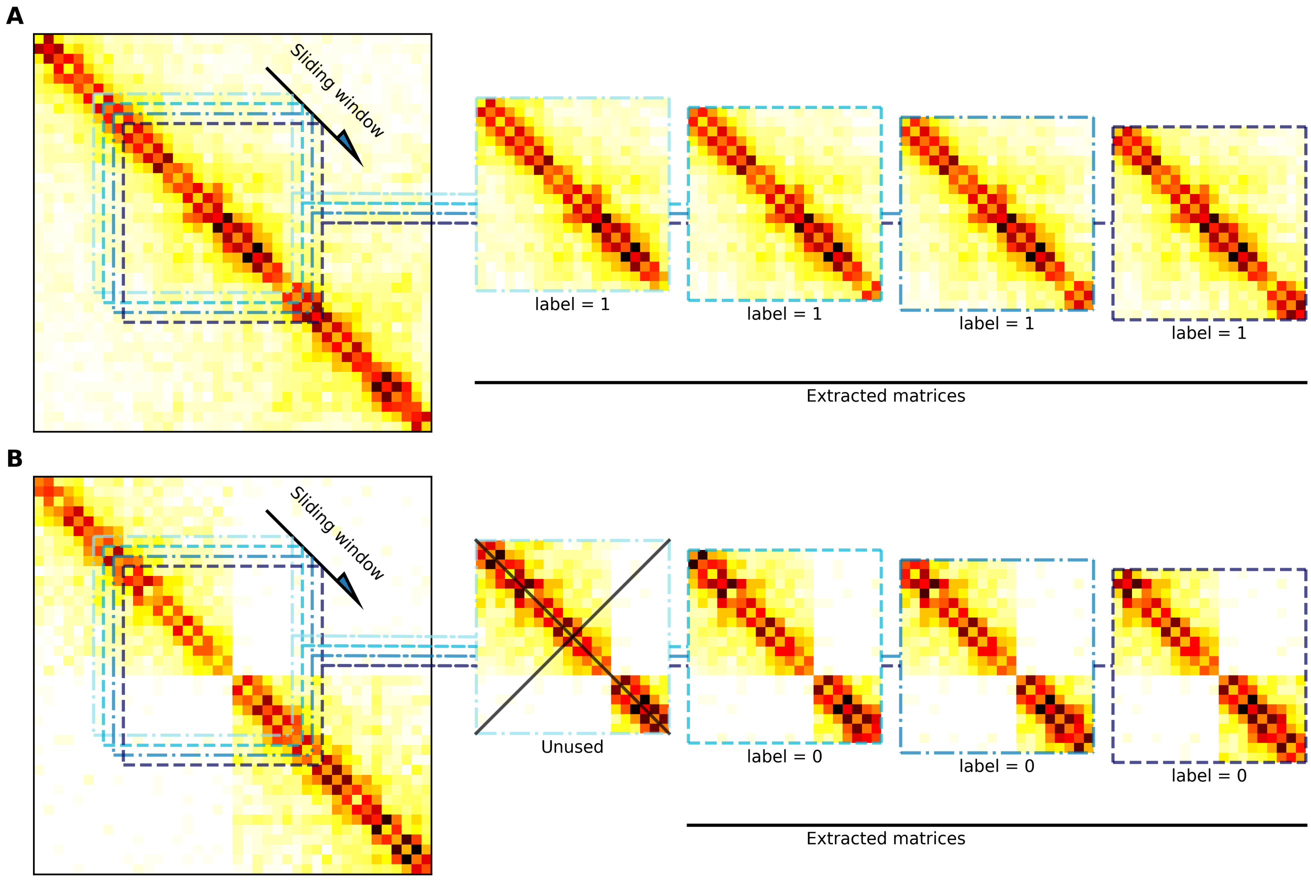


Figure S1: Sample extraction from whole Hi-C matrices using a sliding window along matrices diagonals. Hi-C matrices are at 5 kb bin resolution. (A) "Join" samples extracted from a T2T chromosome. All samples are labeled as 1. (B) "Misjoin" samples extracted from a chimeric contig simulating a deletion. Samples that are close to the breakpoint location (center of the whole matrice) are extracted and labeled as 0.

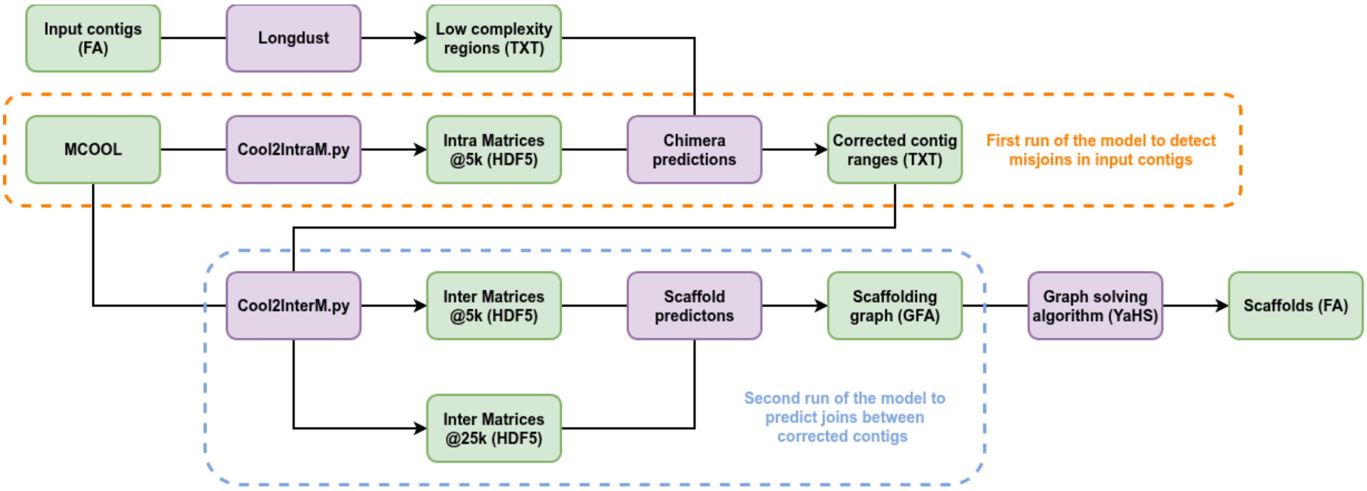


Figure S2: Steps of SPA-C workflow. Input contigs are first screened for misjoins using the *SPA-C* model. Predictions are improved in low complexity regions using *Longdust* predictions. After correction, all possible joins between resulting contigs are evaluated using *SPA-C* model, yielding a scaffold assembly graph (GFA). By default, several Hi-C matrix resolution are used (5kb and 25kb). This graph is finally solved into scaffolds using *YaHS* algorithm only.

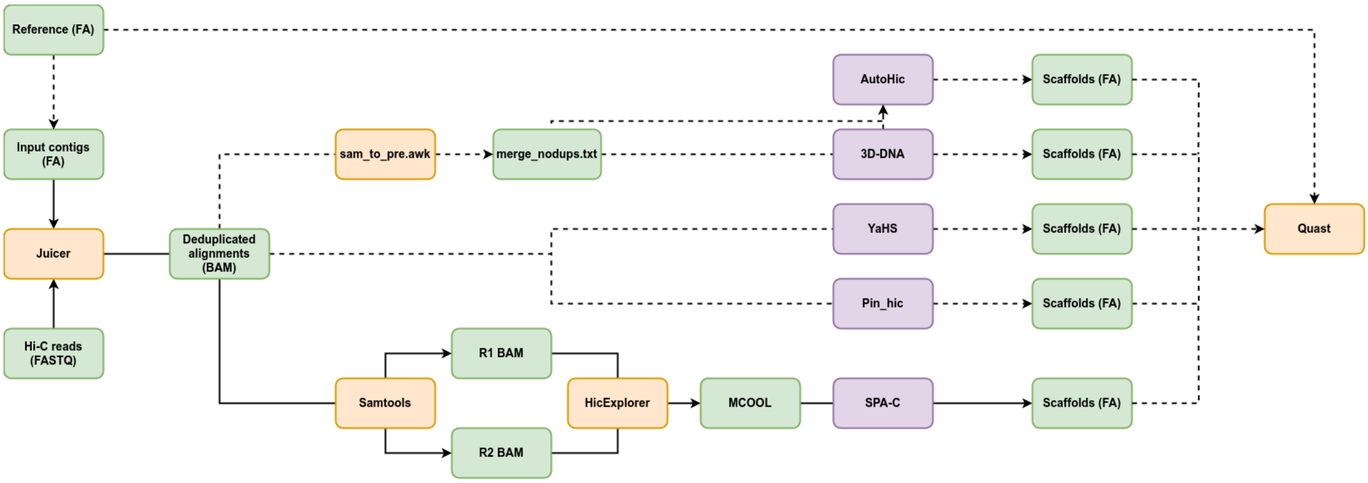


Figure S3: Workflow used to scaffold an input assembly and benchmark scaffolders using Quast. Dashed-lines indicates steps used during benchmarking, solid lines indicate the steps used to run *SPA-C*. For benchmarking only, a reference assembly was broken at each scaffolding gaps into contigs. Hi-C reads are aligned to the input contigs and de-duplicated using *Juicer*. Alignments are passed to *HicExplorer* to generate Hi-C contact matrices in MCOOL format, processed by the *sam_to_pre.awk* script from *Juicer* to be used by *3D-DNA* and *AutoHic*. *YaHS* and *Pin_hic* directly use the deduplicated aligments to scaffold the input assembly. Scaffolds from all tools are processed and compared to the reference using Quast to compute NGA$x$ metrics and the number of misassemblies.

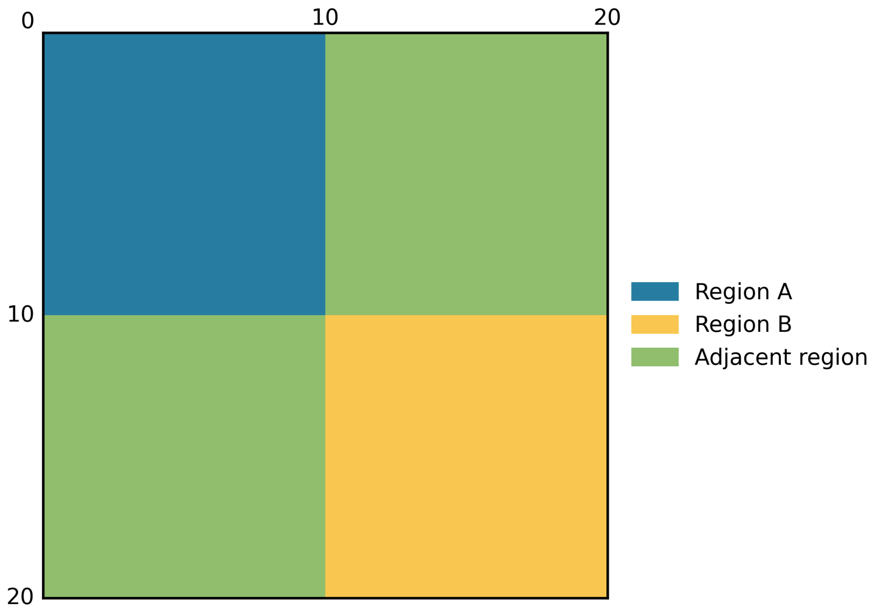


Figure S4: Input structure for the model. The matrix is composed of four quadrants: two “intra” regions and two “inter” or “adjacent” regions. In case of a “join” (correct) sample, the “adjacent” quadrants are “intra” regions.

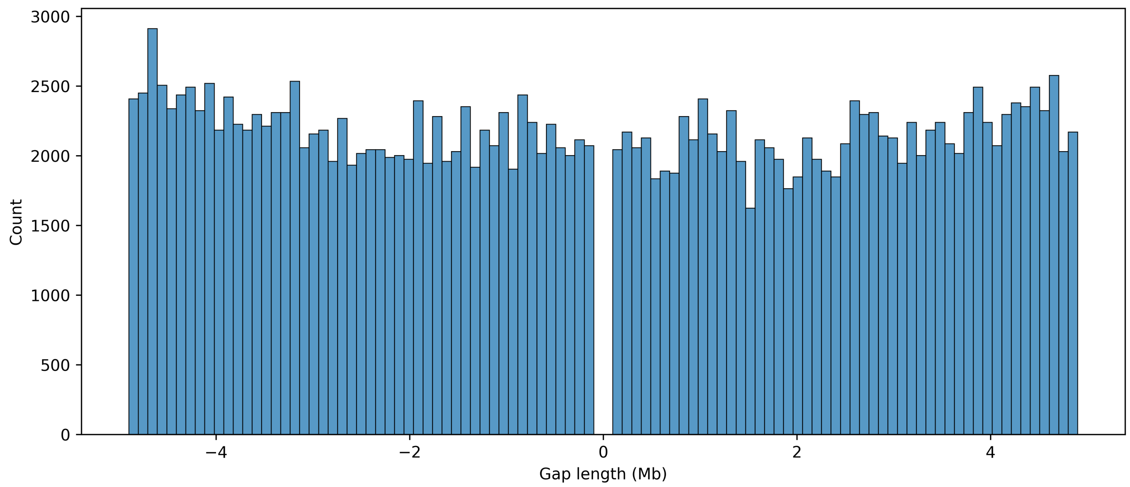


Figure S5: Distribution of gap length between regions of synthetic deletions. Deletions were created by sampling two non-contiguous regions of the same chromosome. A negative gap length indicates that the left region is positioned further along the chromosome than the right region.

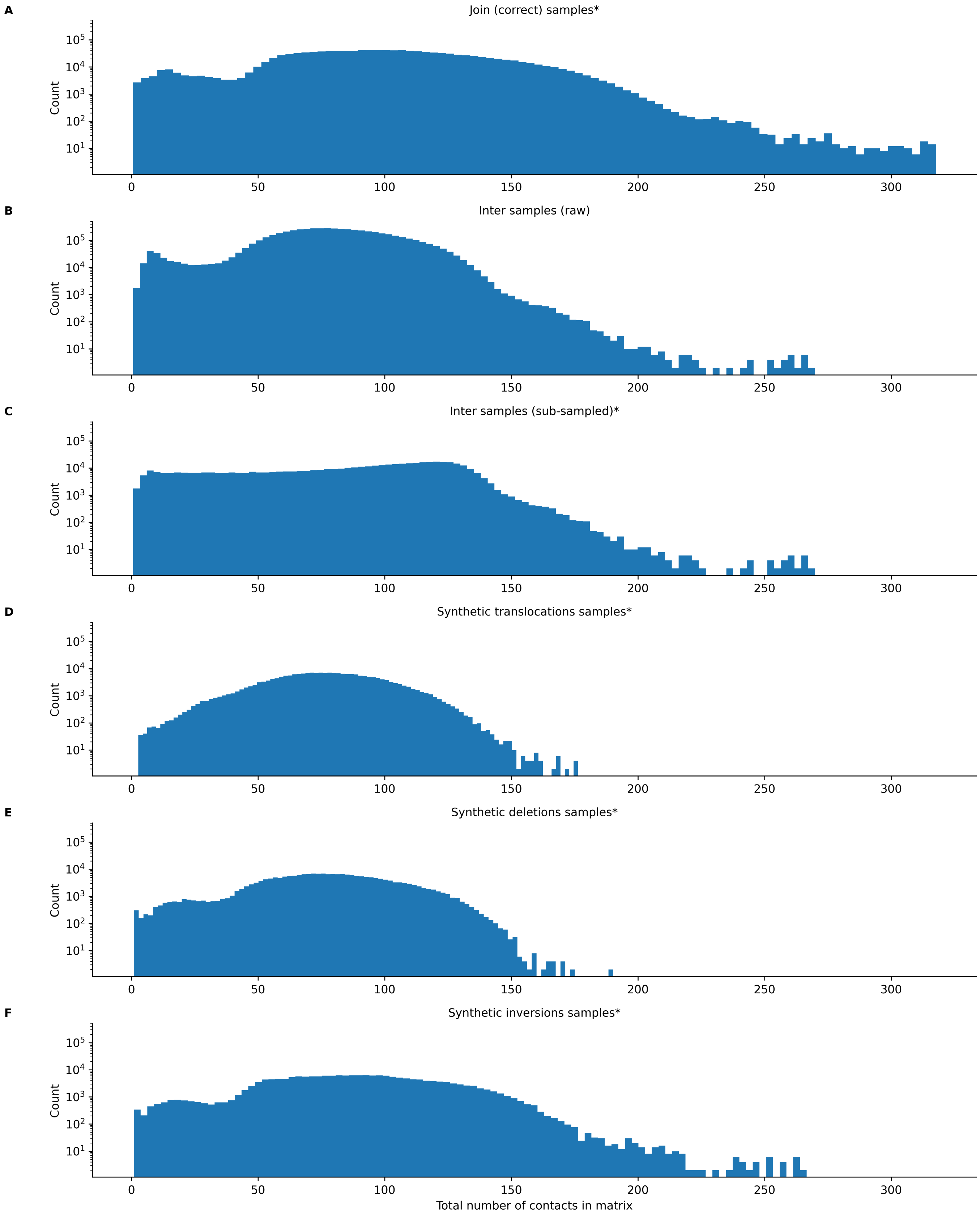


Figure S6: Distribution of Hi-C matrices in subsets of the CHM13v2 dataset by number of contacts. "*" subsets are used in the dataset. Matrices were not normalised. (A) Join samples extracted along the diagonal of chromosomes. (B) Inter samples are misjoin samples extracted from inter-chromosomal regions. Raw distribution (C) Inter samples after sub-sampling of matrices with fewer contacts. (D-F) Misjoin samples extracted along the diagonal of synthetic chimeric contigs representing a type of assembly error.

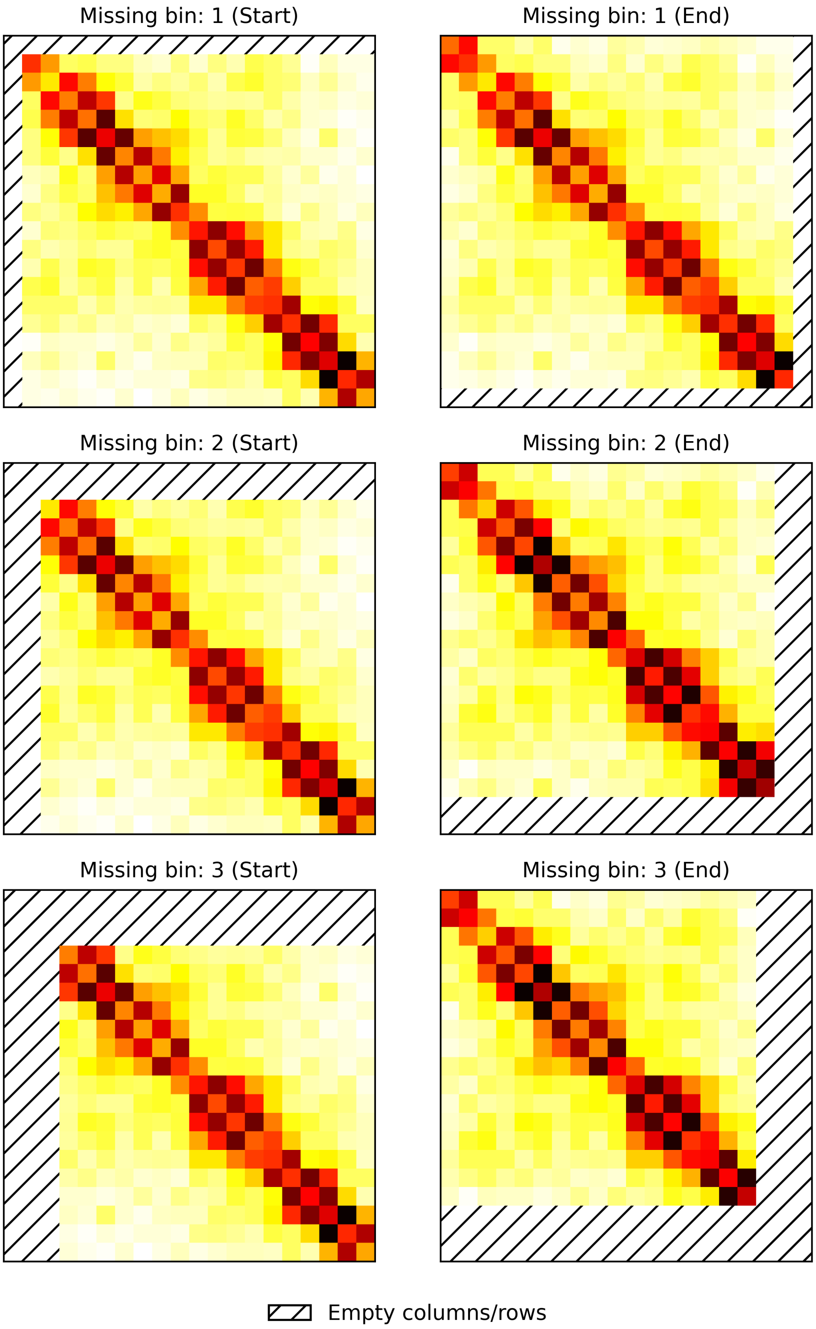


Figure S7: Visualisation of empty bins on an input sample. Bins are always masked from the upper-left (Start) or the lower-right (End) to fill the input required size. Hi-C matrices are at 5 kb bin resolution.

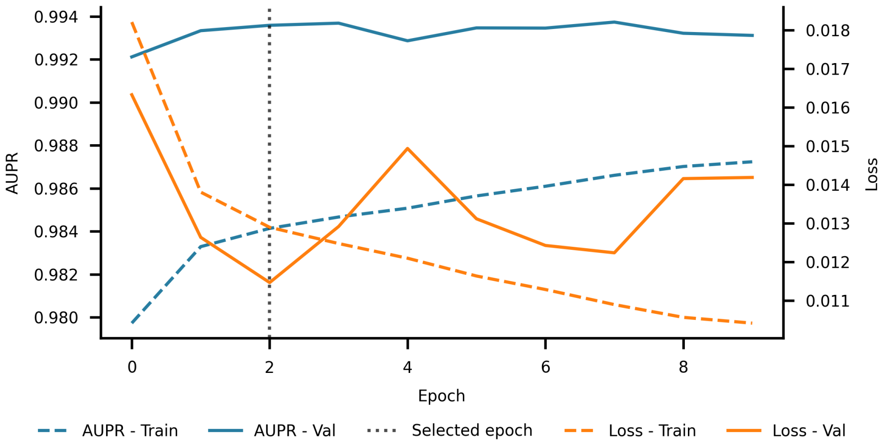


Figure S8: Training curves of the model. Train and Val values are computed using respectively the *train* and *val* subsets of the CHM13v2 dataset. The selected epoch correspond to the version of the weights that are used by the model in the paper.

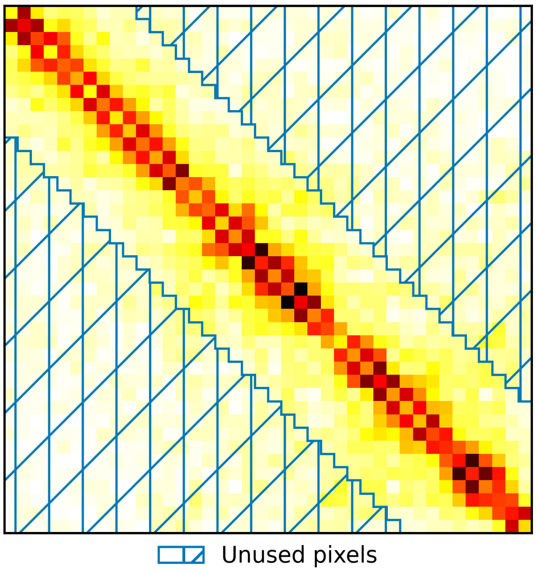


Figure S9: Pixels used in the computation of the 95th quantile. Hi-C matrices are at 5 kb bin resolution. Q95 is then used as maximum value in the minmax rescaling normalisation.

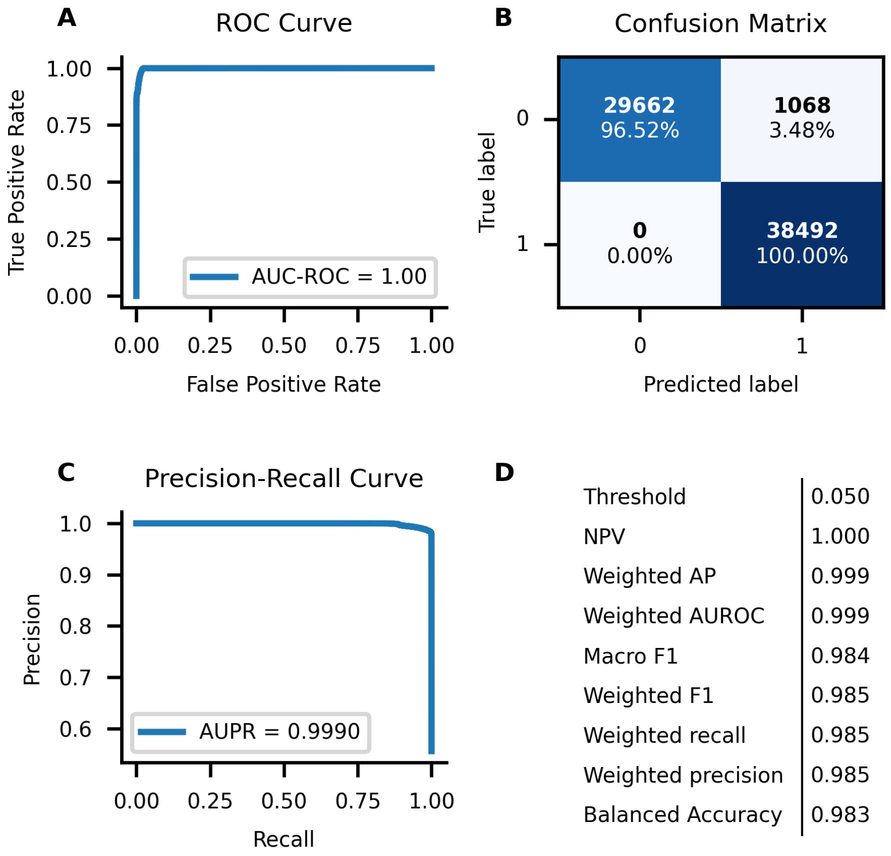


Figure S10: Classification performance of *SPA-C* model on the test subset of the CHM13v2 dataset. (A) ROC Curve and AUC-ROC. (B) Confusion matrix using a threshold of 0.05. Label 0 corresponds to examples of assembly error. (C) PR Curve and AUPR. (D) Several classification performance metrics using a threshold of 0.05.

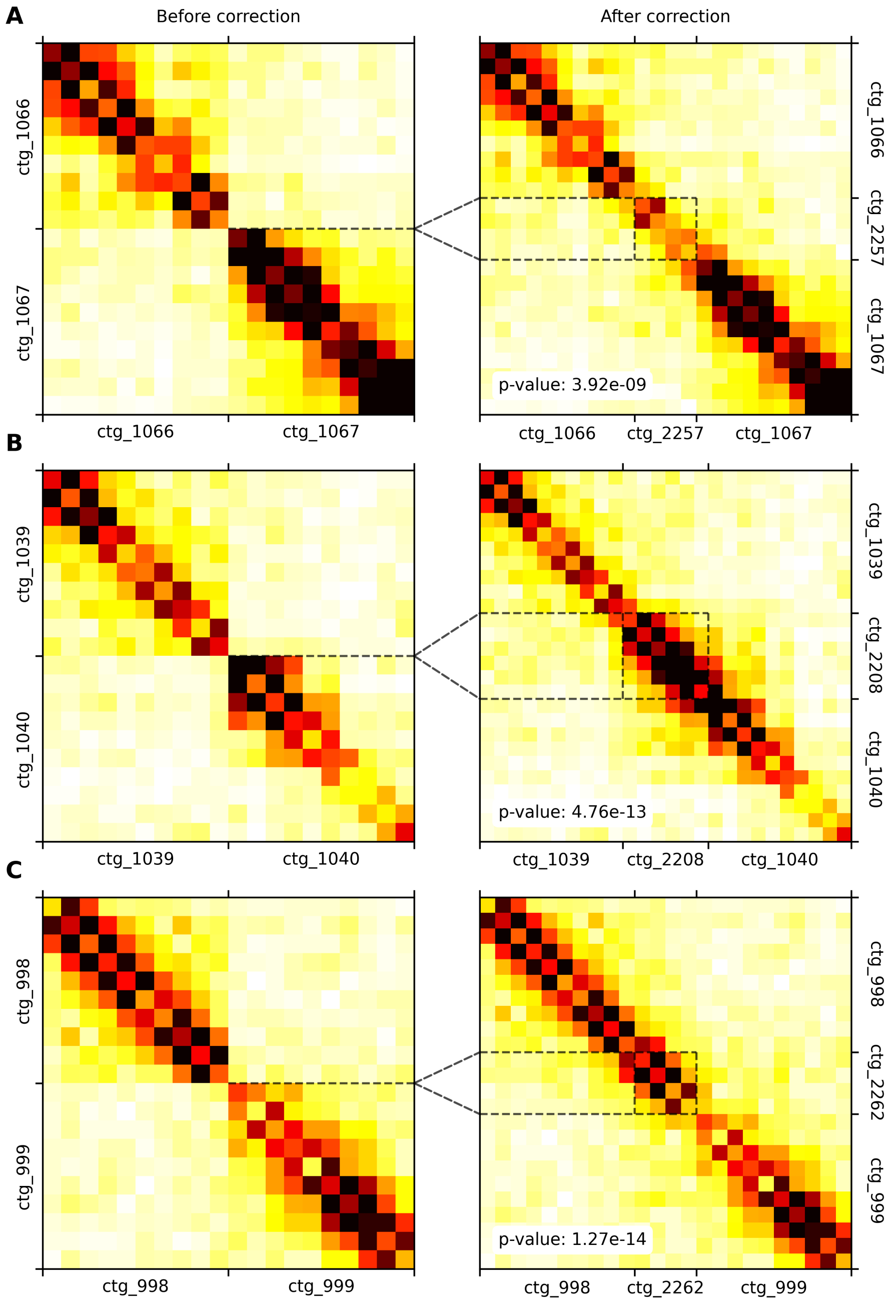


Figure S11: Improvements of the reference assembly made by *SPA-C* on the *V. vulpes* dataset. 5 kb bin resolution matrices are normalised. P-values are associated with the Wilcoxon "greater" alternative test between contact distribution before and after correction. Here, all p-values are significant, indicating that the distribution of contacts is greater after the insertion. Dotted lines show both the insertion position in the left matrix and the inserted contig boundaries in the right panel. (All) Example on the Chromosome 8 of the reference assembly.

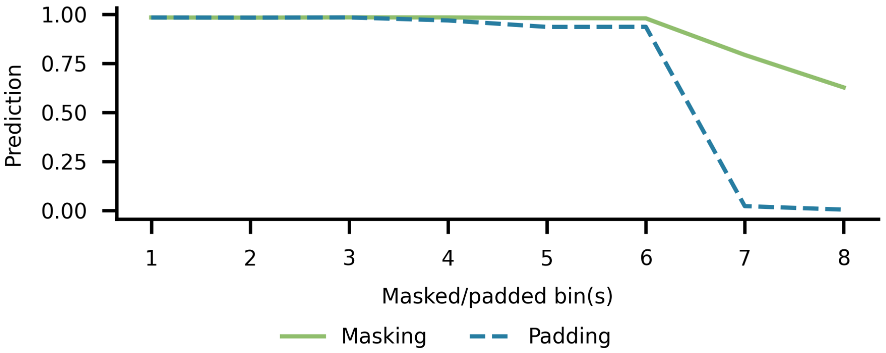


Figure S12: Sensitivity of the *SPA-C* model to empty bins. As the model's input size is fixed to 20 bins or 100 kb, smaller contigs might not be long enough to cover the entire input size, leading to empty bins. We also compared masking through partial convolution with regular padding. Bins were masked or padded starting from the outer edges of the input. Top-left and bottom-right masked/padded bins predictions are aggregated.

#### Supplementary tables

Table S1: Comparison of ResNeXt-50 and *SPA-C*'s model architecture. Since *SPA-C* inputs are much smaller than ResNeXt-50 inputs (20×20 vs 224×224), fewer convolutions are required to decrease the size of the input. Therefore, many ResNeXt blocks were removed resulting in a huge difference in the number of parameters. Within ResNeXt blocks, settings are: filter size, number of channels, cardinality. The repetition of blocks is given outside the array.

| stage | ResNeXt-50 (32×4d) | | SPA-C | |
|---|---|---|---|---|
| | output | layers | layers | output |
| conv1 | 112×112 | 7×7, 64, stride 2<br>3×3 max pool, stride 2 | 7×7, 16, stride 2, padding 3 | 10×10 |
| conv2 | 56×56 | $\begin{bmatrix} 1\times1, 128 \\ 3\times3, 128, C=32 \\ 1\times1, 256 \end{bmatrix} \times 3$ | $\begin{bmatrix} 1\times1, 8 \\ 3\times3, 8, C=4 \\ 1\times1, 32 \end{bmatrix} \times 1$ | 10×10 |
| conv3 | 28×28 | $\begin{bmatrix} 1\times1, 256 \\ 3\times3, 256, C=32 \\ 1\times1, 512 \end{bmatrix} \times 4$ | $\begin{bmatrix} 1\times1, 16 \\ 3\times3, 16, C=4 \\ 1\times1, 64 \end{bmatrix} \times 1$ | 10×10 |
| conv4 | 14×14 | $\begin{bmatrix} 1\times1, 512 \\ 3\times3, 512, C=32 \\ 1\times1, 1024 \end{bmatrix} \times 6$ | $\begin{bmatrix} 1\times1, 16 \\ 3\times3, 16, C=4 \\ 1\times1, 64 \end{bmatrix} \times 1$ | 5×5 |
| conv5 | 7×7 | $\begin{bmatrix} 1\times1, 1024 \\ 3\times3, 1024, C=32 \\ 1\times1, 2048 \end{bmatrix} \times 3$ | | |
| | 1×1 | global average pool<br>1000-d fc, softmax | global average pool<br>32-d fc, 1-d fc, sigmoid | 1×1 |
| # params. | | $25.0\times10^6$ | 42.241 | |

Table S2: Datasets and associated data sources.

| Dataset | Accession | Main link | Assembly download link | Hi-C reads download links |
|---|---|---|---|---|
| CHM13v2 | GCA_009914755.4 | Github | chm13v2.0_noY.fa.gz | R1<br>R2 |
| HG002v1.1 | GCA_018852615.3 | Github | hg002v1.1.fasta.gz | R1<br>R2 |
| *A. gentilis* | GCA_929443795 | portal.darwintreeoflife.org | bAccGen1.2_genomic.fna.gz | R1<br>R2 |
| *A. thaliana* | GCA_933208065 | portal.darwintreeoflife.org | ddAraThal4.1_genomic.fna.gz | R1<br>R2 |
| *B. physalus* | GCA_933208065 | portal.darwintreeoflife.org | mBalPhy2.hap2.1_genomic.fna.gz | R1<br>R2 |
| *D. histrio* | GCA_958299025 | portal.darwintreeoflife.org | idDroHist2.2_genomic.fna.gz | R1<br>R2 |
| *V. vulpes* | GCA_964106825 | portal.darwintreeoflife.org | mVulVul1.hap1.2_genomic.fna.gz | R1<br>R2 |

Table S3: Comparison of *SPA-C* overall performance, with and without *Longdust*. *Longdust* is used to detect low complexity regions from the sequence and discard the model's prediction in such regions. The difference made by using *Longdust* is reported in percentage for each dataset. We used high-quality, curated genomes from the Darwin Tree of Life project and the HG002v1.1 maternal haplotype.

| Dataset | | NGA50[1] | NGA90[1] | auNGA[1] | # misassemblies |
|---|---|---|---|---|---|
| | SPA-C | 154.34 | 78.91 | 153.69 | 3.00 |
| HG002 | SPA-C without Longdust | 143.80 | 66.07 | 146.15 | 6.00 |
| | Difference | 7.33% | 19.43% | 5.16% | -50.00% |
| | SPA-C | 27.69 | 15.78 | 25.19 | 1.00 |
| *A. thaliana* | SPA-C without Longdust | 14.46 | 1.93 | 12.23 | 6.00 |
| | Difference | 91.50% | 719.54% | 105.97% | -83.33% |
| | SPA-C | 23.34 | 0.81 | 20.23 | 32.00 |
| *D. histrio* | SPA-C without Longdust | 23.34 | 0.55 | 20.21 | 36.00 |
| | Difference | 0.00% | 48.09% | 0.11% | -11.11% |
| | SPA-C | 21.43 | 2.61 | 22.52 | 124.00 |
| *A. gentilis* | SPA-C without Longdust | 21.03 | 0.85 | 21.34 | 180.00 |
| | Difference | 1.90% | 208.71% | 5.52% | -31.11% |
| | SPA-C | 35.80 | 7.79 | 39.96 | 193.00 |
| *V. vulpes* | SPA-C without Longdust | 34.69 | 7.65 | 36.23 | 187.00 |
| | Difference | 3.22% | 1.82% | 10.28% | 3.21% |
| | SPA-C | 30.05 | 0.35 | 34.86 | 468.00 |
| *B. physalus* | SPA-C without Longdust | 26.34 | 0.31 | 33.23 | 487.00 |
| | Difference | 14.10% | 12.90% | 4.90% | -3.90% |

[1] Expressed in Mbp.

Table S4: Evaluation of *SPA-C* model's performance on other bin resolutions. The model was trained using 5 kb bin resolution Hi-C matrices. The dataset used in this table was built using the maternal haplotype of HG002v1.1 T2T assembly. Chimeric contigs were built using 2Mb-size regions instead of the 100kb size regions of the training dataset to allow for lower bin resolution.

| Bin resolution | AUPR | macro F1 | Number of samples | Proportion of correct samples (%) |
|---|---|---|---|---|
| 5k | 0.9985 | 0.9898 | 1,712,136 | 71.24 |
| 10k | 0.9992 | 0.9932 | 946,938 | 64.36 |
| 25k | 0.9960 | 0.9939 | 657,610 | 36.99 |
| 100k | 0.9989 | 0.9861 | 99,552 | 60.40 |

Table S5: Comparison of *SPA-C* overall performance, with and without multi-resolution. With multi-resolution, the model predicts linkage pseudo-probability for each bin resolution independently. Similar predictions are then averaged between resolutions. We compared a default *SPA-C* run which uses 5 kb and 25 kb bin resolution matrices with a run using 5 kb only. The difference made by using several resolutions is reported in percentage for each dataset. We used high-quality, curated genomes from the Darwin Tree of Life project and the HG002v1.1 maternal haplotype.

| Dataset | | NGA50[1] | NGA90[1] | auNGA[1] | # misassemblies |
|---|---|---|---|---|---|
| | SPA-C | 154.34 | 78.91 | 153.69 | 3.00 |
| HG002 | SPA-C@5k | 154.34 | 78.91 | 153.69 | 1.00 |
| | Difference | 0.00% | 0.00% | 0.00% | 200.00% |
| | SPA-C | 27.69 | 15.78 | 25.19 | 1.00 |
| *A. thaliana* | SPA-C@5k | 27.69 | 19.63 | 26.84 | 5.00 |
| | Difference | 0.00% | -19.63% | -6.15% | -80.00% |
| | SPA-C | 23.34 | 0.81 | 20.23 | 32.00 |
| *D. histrio* | SPA-C@5k | 21.02 | 0.69 | 17.39 | 30.00 |
| | Difference | 11.07% | 17.20% | 16.30% | 6.67% |
| | SPA-C | 21.43 | 2.61 | 22.52 | 124.00 |
| *A. gentilis* | SPA-C@5k | 16.46 | 1.60 | 18.16 | 131.00 |
| | Difference | 30.19% | 62.74% | 24.03% | -5.34% |
| | SPA-C | 35.80 | 7.79 | 39.96 | 193.00 |
| *V. vulpes* | SPA-C@5k | 36.31 | 6.76 | 36.32 | 167.00 |
| | Difference | -1.39% | 15.17% | 10.01% | 15.57% |
| | SPA-C | 30.05 | 0.35 | 34.86 | 468.00 |
| *B. physalus* | SPA-C@5k | 20.01 | 0.35 | 27.19 | 481.00 |
| | Difference | 50.16% | 0.00% | 28.20% | -2.70% |

[1] Expressed in Mbp.

Table S6: CPU time of every steps of the benchmarking section. Since the number of threads is not equal between tasks, the cumulative wall clock is not reported.

| | Dataset | Juicer | | HicExplorer | | Merge_nodups[1] | | Scaffolder | | CPU time | |
|---|---|---|---|---|---|---|---|---|---|---|---|
| | | time | #threads | time | #threads | time | #threads | time | #threads | sec. | hours |
| YaHS | HG002 | 31:13:17 | 16 | - | - | - | - | 01:13:31 | 4 | 1815996 | 504 |
| | *A. thaliana* | 12:56:33 | 16 | - | - | - | - | 25:50:00 | 4 | 1117488 | 310 |
| | *D. histrio* | 14:03:20 | 16 | - | - | - | - | 00:30:03 | 4 | 816812 | 226 |
| | *A. gentilis* | 10:22:09 | 16 | - | - | - | - | 00:20:39 | 4 | 602220 | 167 |
| | *V. vulpes* | 28:10:54 | 16 | - | - | - | - | 01:34:23 | 16 | 1713872 | 476 |
| | *B. physalus* | 34:51:21 | 24 | - | - | - | - | 01:36:34 | 4 | 3034720 | 842 |
| Pin_hic | HG002 | 31:13:17 | 16 | - | - | - | - | 03:34:11 | 4 | 1849756 | 513 |
| | *A. thaliana* | 12:56:33 | 16 | - | - | - | - | 01:21:21 | 4 | 765012 | 212 |
| | *D. histrio* | 14:03:20 | 16 | - | - | - | - | 01:36:33 | 4 | 832772 | 231 |
| | *A. gentilis* | 10:22:09 | 16 | - | - | - | - | 00:51:15 | 4 | 609564 | 169 |
| | *V. vulpes* | 28:10:54 | 16 | - | - | - | - | 04:19:11 | 16 | 1872080 | 520 |
| | *B. physalus* | 34:51:21 | 24 | - | - | - | - | 04:27:49 | 4 | 3075820 | 854 |
| 3D-DNA | HG002 | 31:13:17 | 16 | - | - | 02:21:05 | 8 | 62:58:43 | 4 | 2772964 | 770 |
| | *A. thaliana* | 12:56:33 | 16 | - | - | 01:04:25 | 8 | 16:49:45 | 1 | 836993 | 232 |
| | *D. histrio* | 14:03:20 | 16 | - | - | 01:13:50 | 8 | 11:52:40 | 1 | 887800 | 246 |
| | *A. gentilis* | 10:22:09 | 16 | - | - | 00:58:05 | 8 | 44:12:09 | 1 | 784273 | 217 |
| | *V. vulpes* | 28:10:54 | 16 | - | - | 05:11:11 | 8 | 51:50:28 | 16 | 4758680 | 1321 |
| | *B. physalus* | 34:51:21 | 24 | - | - | 04:45:51 | 8 | 71:13:41 | 4 | 4174436 | 1159 |
| AutoHic | HG002 | 31:13:17 | 16 | - | - | 02:21:05 | 8 | 41:21:47 | 4 | 3368592 | 935 |
| | *A. thaliana* | 12:56:33 | 16 | - | - | 01:04:25 | 8 | 01:39:09 | 4 | 860789 | 239 |
| | *D. histrio* | 14:03:20 | 16 | - | - | 01:13:50 | 8 | 01:12:52 | 4 | 905288 | 251 |
| | *A. gentilis* | 10:22:09 | 16 | - | - | 00:58:05 | 8 | 25:23:39 | 4 | 1149949 | 319 |
| | *V. vulpes* | 28:10:54 | 16 | - | - | 05:11:11 | 8 | - | - | 4758680 | 1321 |
| | *B. physalus* | 34:51:21 | 24 | - | - | 04:45:51 | 8 | - | - | 4174436 | 1159 |
| SPA-C | HG002 | 31:13:17 | 16 | 10:13:18 | 4 | - | - | 00:17:33 | 4 | 1949756 | 541 |
| | *A. thaliana* | 12:56:33 | 16 | 02:51:05 | 4 | - | - | 00:02:09 | 4 | 787064 | 218 |
| | *D. histrio* | 14:03:20 | 16 | 02:51:24 | 4 | - | - | 00:07:03 | 4 | 852428 | 236 |
| | *A. gentilis* | 10:22:09 | 16 | 01:58:03 | 4 | - | - | 00:20:20 | 4 | 630476 | 175 |
| | *V. vulpes* | 28:10:54 | 16 | 13:06:06 | 4 | - | - | 02:49:30 | 4 | 1852608 | 514 |
| | *B. physalus* | 34:51:21 | 24 | 09:40:52 | 8 | - | - | 07:37:12 | 4 | 3400088 | 944 |

[1] Conversion of merged_dedup.bam file to merged_nodups.txt using sam_to_pre.awk script from *Juicer*.

### Supplementary methods

#### Hi-C read alignments and matrix generation

We used *Samtools* (v1.21) to index genomes and work with BAM files. We used the *Juicer* (v2, commit 3e2ab76) workflow to align and de-duplicate Hi-C reads using default parameters and the *-e* (early exit) setting. This stopped the workflow at the generation of the *merged_dedup.bam* file. As this BAM contains both mates of each paired reads, we split it into two dedicated R1 and R2 BAMs using *samtools view -b -f 64* and *samtools view -b -f 128* respectively. R1 and R2 BAMs were then passed to the *hicBuildMatrix* (v3.7.6) tool from *HicExplorer* using –*skipDuplicationCheck –minMappingQuality 0* arguments, creating a MCOOL file. Scripts are available at `https://github.com/SPA-C/SPA-C`.

#### MCOOL file format

MCOOL files are based on the well-described HDF5 file format. See `https://cooler.readthedocs.io/en/latest/schema.html` for more information.

#### *SPA-C* inference workflow and implementation

Given an input assembly (FASTA), a "intra" dataset is generated using the *Cool2Intra_FM.py* script from the MCOOL file. It extracts matrices along contig's diagonals into an HDF5 file, for contigs of length $\geq$ 200 kb only. *Longdust* is then run on the input assembly. The *Chimera_predictor.py* script uses the "intra" dataset and *Longdust* predictions to predict scores along contigs, detects misjoins and outputs a list of contig ranges that are free of assembly errors in a simple text file. The script can be run without *Longdust* predictions as well. The *Cool2Inter_FM.py* script takes error-free contig ranges and the original MCOOL file to generate "inter" dataset. The script is run for each bin resolution (5 and 25 kb by default), yielding two HDF5 files. These files, as well as the input FASTA are passed to the *Scaffold_predictor.py* script in which linkage score are predicted by the model, aggregated in bin resolution-specific graphs, then concatenated into a final scaffolding graph and output as a GFA file. This GFA is finally passed to the scaffolding algorithm of *YaHS*, creating the final assembly (FASTA) and an AGP file describing the choices made by the algorithm. We containerised environments needed to run *SPA-C* into two separate apptainer images due to incompatibility issues. One contains *HicExplorer*, *Cooler* and their dependencies. The second is based on the *PyTorch* GPU-enabled docker image and used to run the model. Both containers made it simple to use *SPA-C* on any system. All steps are regrouped into a one-click bash script for easy use. Parameters remains transparent to the user within the bash script. Scripts are available at `https://github.com/SPA-C/SPA-C` and Apptainer images repository are available at `https://github.com/SPA-C/ENV` and `https://github.com/SPA-C/HiC-ENV`.

#### Hardware and computing ressources

Training and testing of the model was done using an NVIDIA RTX 3050 GPU. We used a CPU cluster to benchmark scaffolders. CPU time is given as the sum of *number of allocated threads*$\times$*job wall time* in Tab.1, for each necessary steps. As we did not allocate entire nodes and some tools/steps used different CPU/memory settings, CPU times given in Tab.1 should be used as an indication rather than a precise measure of each tool computation speed. The full table is also available in Tab.S
